# Ability of known colorectal cancer susceptibility SNPs to predict colorectal cancer risk: A cohort study within the UK Biobank

**DOI:** 10.1101/2021.04.28.441750

**Authors:** Aviv Gafni, Gillian S. Dite, Erika Spaeth Tuff, Richard Allman, John L. Hopper

## Abstract

Colorectal cancer risk stratification is crucial to improve screening and risk-reducing recommendations, and consequently do better than a one-size-fits-all screening regimen. Current screening guidelines in the UK, USA and Australia focus solely on family history and age for risk prediction, even though the vast majority of the population do not have any family history. We investigated adding a polygenic risk score based on 45 single-nucleotide polymorphisms to a family history model (combined model) to quantify how it improves the stratification and discriminatory performance of 10-year risk and full lifetime risk using a prospective population-based cohort within the UK Biobank. For both 10-year and full lifetime risk, the combined model had a wider risk distribution compared with family history alone, resulting in improved risk stratification of nearly 2-fold between the top and bottom risk quintiles of the full lifetime risk model. Importantly, the combined model can identify people (n=72,019) who do not have family history of colorectal cancer but have a predicted risk that is equivalent to having at least one affected first-degree relative (n=44,950). We also confirmed previous findings by showing that the combined full lifetime risk model significantly improves discriminatory accuracy compared with a simple family history model 0.673 (95% CI 0.664–0.682 versus 0.666 (95% CI 0.657–0.675), p=0.0065. Therefore, a combined polygenic risk score and first-degree family history model could be used to improve risk stratified population screening programs.

## Introduction

Colorectal cancer is the fourth deadliest cancer, causing nearly 900,000 deaths every year globally. Worldwide, colorectal cancer is the 2nd most common cancer in women and the 3rd in men, with men having around 25% higher incidence and mortality compared with women (1, 2). Colorectal cancer has several non-modifiable risk factors, including age, family history, sex and genetic makeup. Roughly 5%–10% of colorectal cancer cases have an affected first-degree relative, and the strength of associated risk depends on the number and closeness of the relationship, and on the ages at diagnosis of the affected relative(s) and the age of the at risk consult (3–9). Efforts to better understand heritability of colorectal cancer in family studies underscore the complex relationship with environmental components (10, 11).

Rare high penetrance mutations have been found to cause hereditary colorectal cancers, including those predisposing to Lynch syndrome and familial adenomatous polyposis, accounting for 5%–7% of all colorectal cancer cases. Known genetic mutations account for only half of the cases in persons with such family histories (12). The unexplained causes of cases with a family history could be due to polygenic factors, such as common low penetrance single-nucleotide polymorphisms (SNPs) (13, 14) or lifestyle causes that are also shared by family members (15).

In recent years, an increasing number of susceptibility SNPs have been identified by genome-wide association studies, which examine vast numbers of variants across the genome for associations with disease risk (16, 17). Although each susceptibility SNP has a weak association with colorectal cancer risk, the cumulative association of many SNPs combined as a polygenic risk score (PRS) can result in a substantial risk gradient (in both directions) and is potentially an effective risk stratification method (14, 18). For example, Jenkins et al. (19) used a cohort enriched for family history to show the value of a PRS in stratifying individuals by risk, particularly those with a family history but not found to be carriers of mutations associated with Lynch syndrome or familial adenomatous polyposis. Importantly, their study confirmed that a 45 SNP panel in conjunction with having a family history of colorectal cancer could identify non-trivial proportions of the population who would likely benefit from earlier screening. The use of polygenic risk models to inform targeted screening has potential benefit in clinical genetics settings for families in which high-risk mutations cannot be identified (18). Notwithstanding that observation, the reality is that about 90% of colorectal cancer cases have no family history in first-degree relatives and it is this group that could benefit from improved risk prediction (3). Given the incidence of colorectal cancer diagnosed before age 50 years is increasing (20, 21), it is particularly important to focus on risk prediction to accurately identify at-risk adults who may not be identified by current standard screening guidelines. Therefore, there is an important justification for improved risk prediction tools to guide screening and risk reduction.

Our aim is to investigate whether better risk stratification can be achieved in the general population, using the UK Biobank, a prospective population-based cohort. To this end, we have investigated the ability of a model comprising 45 SNPs (PRS) and first-degree family history to stratify risk in the general population and the discriminatory performance and calibration of the model to inform the potential utility in broad application risk stratified screening.

## Methods

### Study sample

The UK Biobank is a major biomedical database, comprises of 500,000 volunteers who were aged 40–69 years when recruited in 2006–2010 from England, Scotland and Wales. The purpose of the UK Biobank is to assist researchers in studying disease prevention, diagnosis and treatment and investigate the determinants of a wide spectrum of diseases in middle and later life (22, 23). The UK Biobank has Research Tissue Bank approval (REC #16/NW/0274) that covers analysis of data by approved researchers. All participants provided written informed consent to the UK Biobank before data collection began. This research has been conducted using the UK Biobank resource under Application Number 47401.

Each participant has provided detailed personal and medical history information and has undergone physical and biological measurements. Samples provided include blood, urine and saliva. All participants who provided blood have been genotyped and genome-wide SNP data is available for each (24). All participants have agreed to their health status being followed-up via linkage to health registries and general practice and hospital records. Therefore, the UK Biobank is a powerful resource to study genetic associations and disease risk due to being a prospective cohort, its large size, and the wealth of genetic and clinical information it has and will collect. The eligibility criteria for this study are described in Table 1.

**Table 1:** Eligibility criteria.

| **N eligible** | **Criteria** | **N dropped** |
| --- | --- | --- |
| 502,488 | Active participants in UK Biobank |  |
| 487,869 | Reported sex same as genetically determined sex | 14,619 |
| 409,289 | White British and genetically Caucasian | 78,580 |
| 406,745 | No previous diagnosis of colorectal cancer at baseline | 2,544 |
| 404,715 | Aged 40–69 years at assessment date | 2,030 |
| **403,998** | Genome-wide SNP data available | 717 |

### Generation of PRS

A PRS was calculated for each UK Biobank participant using the 45 SNPs (S1 Table) that were found to be associated with colorectal cancer by previous studies (13, 25). For each SNP, the previously published odds ratio (OR) per risk allele and risk allele frequency (*p*) were used to calculate the population average risk using the formula: μ = (1 − *p*)^2^ + 2*p*(1 − *p*)OR + *p*^2^OR^2^ (26). The population average risk was normalised to 1 using weighted risk values, which were calculated as 1/μ, OR/μ and OR^2^/μ for the three genotypes (defined by the number of risk alleles 0, 1, or 2). The PRS risk score for each participant was calculated by multiplying the weighted risk values for each of the 45 SNPs (assuming independent and additive risks on the log odds scale) (19).

### Outcome

The outcome of interest was invasive colorectal cancer diagnosis after baseline assessment. Colorectal cancer was identified using linked cancer registry data using ICD-9 (1530–1539, 1540–1541), ICD-10 (C18–C20) codes or self-reported disease. Follow-up began at date of baseline assessment and observations were censored at the earliest of date of diagnosis, date of death or 31 March 2016 (the latest date for which linkage to cancer registries is complete), whichever occurred first. For analysis of standardised incidence ratios (SIR) for 10-years of follow-up, we ceased follow-up after 10 years.

### Risk scores

We evaluated the following two models involving: (i) family history only (based on number of affected first-degree relatives) and (ii) a combination of both family history and the PRS (combined model). Relative risks for having 0, 1 or ≥2 first-degree relatives diagnosed with colorectal cancer were obtained from a previous study (27), and centred to have a population average of 1. The PRS model used the 45 SNPs described by Jenkins et al (19). For SNPs rs10904849, rs35509282, rs4925386 and rs10911251, we used surrogate SNPs rs10904850, rs11100443, rs11204472 and rs6669796 respectively, and for 19qhap (19q13.2) and 11qhap (11q12.2), we used the tag SNPs rs1800469 and rs174537 respectively (S1 Table).

Calculation of absolute 10-year risk was performed using sex- and age-specific incidence rates for England in 2013, and took into account competing mortality, obtained from the UK Office for National Statistics (28). For the calculation of the absolute full lifetime risk to age 85, mortality rates were excluded. Risk scores were centred to have a mean of 1. SIRs were calculated using the observed vs expected colorectal cancer incidence based on population gender- and age-specific incidence rates for England in 2006–2016 (29).

### Statistical analysis

#### Model performance

Model discrimination was determined using the area under the receiver operating characteristic curve (AUC). We assessed model calibration using logistic regression analysis, for which the observed colorectal cancer case status was the dependent variable and the log-odds of our model’s predicted probability for the outcome of colorectal cancer during the follow-up time was the independent variable. The test for dispersion was performed by evaluating the null hypothesis that the estimated regression coefficient was equal to 1 in the model without a constant term (30). Overdispersion (regression coefficient <1) occurs when the observed values have greater variability then the expected values produced by the model, while under-dispersion (regression coefficient >1) happens when the observed values show less variation than expected.

Broad sense calibration was measured using 10-year follow-up data from the UK Biobank, for which the SIR (observed/expected incidence) was calculated for both models.

All statistical analyses were performed using Stata version 16.1 (31). All statistical tests were two sided and p < 0.05 was considered nominally statistically significant.

## Results

Characteristics of participants and the mean 10-year and full lifetime risks for the combined model are summarised in Table 2. The mean age at baseline of colorectal cancer cases and controls was 61.45 years (SD 6.33) and 57.28 years (SD 7.96), respectively.

**Table 2:** Summary statistics for the eligible UK biobank cohort.

|  | Unaffected<br>Total 401,006 | Affected (incident<br>cases)<br>Total 2,992 |
| --- | --- | --- |
|  | N (%) | N (%) |
| **Age at cohort entry (years)** |  |  |
| 40–49 | 88,648 (22.11) | 198 (6.62) |
| 50–59 | 133,056 (33.18) | 800 (26.74) |
| 60–69 | 179,302 (44.71) | 1,994 (66.64) |
| **Age when diagnosed with colorectal**<br>**cancer (years)** |  |  |
| 40–49 | – | 79 (2.64) |
| 50–59 | – | 496 (16.58) |
| 60–69 | – | 1,647 (55.05) |
| 70–79 | – | 770 (25.74) |
| **Gender** |  |  |
| Female | 217,501 (54.24) | 1,275 (42.61) |
| Male | 183,505 (45.76) | 1,717 (57.39) |
| **Number of first-degree relatives**<br>**diagnosed with colorectal cancer** |  |  |
| 0 | 356,437 (88.89) | 2,544 (85.03) |
| 1 | 42,129 (10.51) | 412 (13.77) |
| 2+ | 2440 (0.61) | 36 (1.20) |
|  | **Mean (SD)** | **Mean (SD)** |
| **Full lifetime risk: combined model** | 0.068 (0.041) | 0.080 (0.050) |
| **10-year risk: combined model** | 0.011 (0.009) | 0.016 (0.013) |

Overall, the SIR of observed colorectal cancer compared with the number expected using population incidences was 0.92 (95% CI = 0.88–0.95) (Table 3), meaning that the colorectal cancer incidence in the UK Biobank data was 8% (95% CI = 5-12%) less than expected. Furthermore, the SIR broken down by gender showed that the expected incidence for females and males was 6% (95% CI = 1-11%) and 10% (95% CI = 6-14%), respectively, less than expected (Table 3). When the SIR was broken down by age, for ages 60–69 (the majority of cases), the colorectal cancer incidence was ~11% (95% CI = 7-15%) less than expected. This is consistent with the recognized healthy volunteer selection bias of the UK Biobank (32, 33). For ages 50–59 the incidence was ~3% less than expected, and for ages 40–49, the incidence was 6% higher than expected, but the confidence intervals included 1.

**Table 3:**
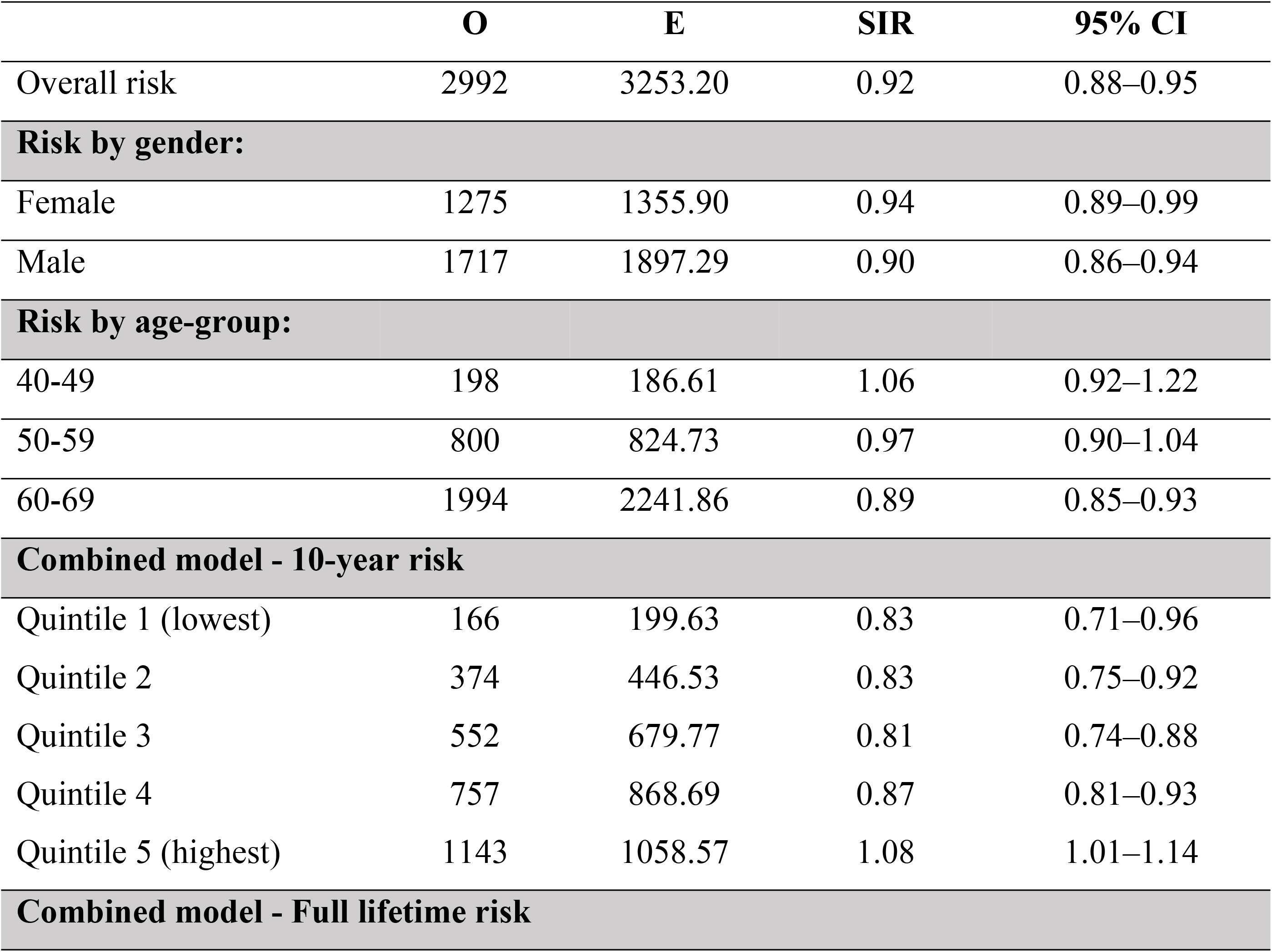
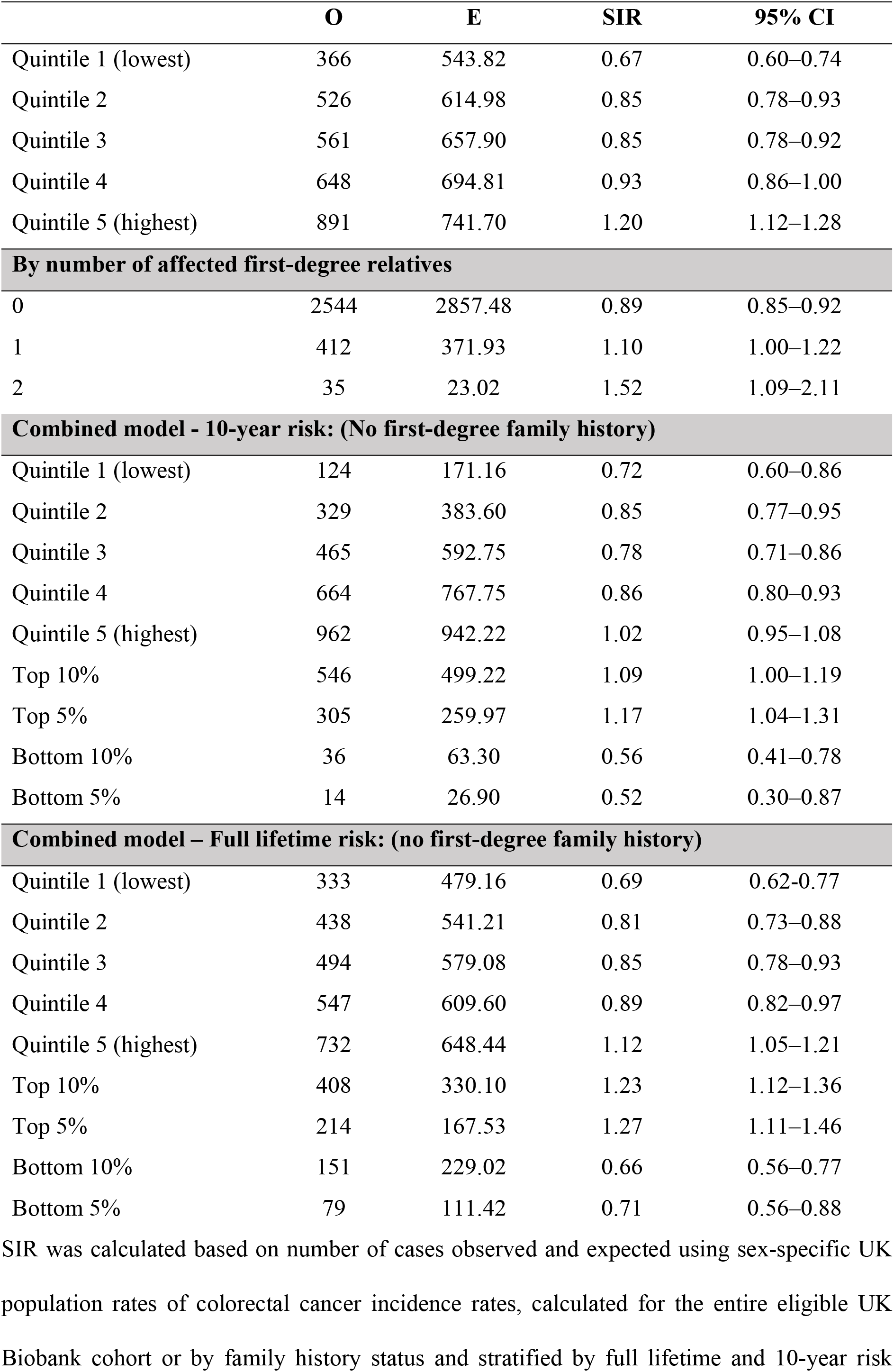
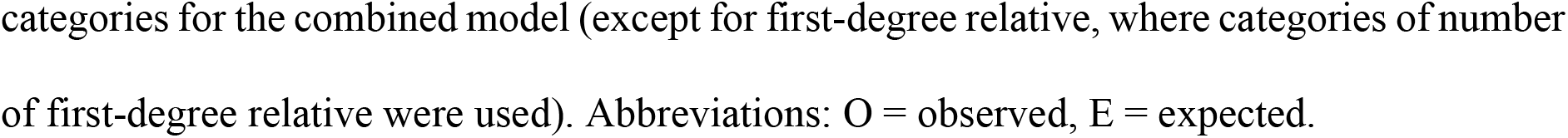
Standardised incidence ratios (SIR) - Overall and by subgroups.

|  | **O** | **E** | **SIR** | **95% CI** |
| --- | --- | --- | --- | --- |
| Overall risk | 2992 | 3253.20 | 0.92 | 0.88–0.95 |
| **Risk by gender:** |  |  |  |  |
| Female | 1275 | 1355.90 | 0.94 | 0.89–0.99 |
| Male | 1717 | 1897.29 | 0.90 | 0.86–0.94 |
| **Risk by age-group:** |  |  |  |  |
| 40-49 | 198 | 186.61 | 1.06 | 0.92–1.22 |
| 50-59 | 800 | 824.73 | 0.97 | 0.90–1.04 |
| 60-69 | 1994 | 2241.86 | 0.89 | 0.85–0.93 |
| **Combined model - 10-year risk** |  |  |  |  |
| Quintile 1 (lowest) | 166 | 199.63 | 0.83 | 0.71–0.96 |
| Quintile 2 | 374 | 446.53 | 0.83 | 0.75–0.92 |
| Quintile 3 | 552 | 679.77 | 0.81 | 0.74–0.88 |
| Quintile 4 | 757 | 868.69 | 0.87 | 0.81–0.93 |
| Quintile 5 (highest) | 1143 | 1058.57 | 1.08 | 1.01–1.14 |
| **Combined model - Full lifetime risk** |  |  |  |  |
| Quintile 1 (lowest) | 366 | 543.82 | 0.67 | 0.60–0.74 |
| Quintile 2 | 526 | 614.98 | 0.85 | 0.78–0.93 |
| Quintile 3 | 561 | 657.90 | 0.85 | 0.78–0.92 |
| Quintile 4 | 648 | 694.81 | 0.93 | 0.86–1.00 |
| Quintile 5 (highest) | 891 | 741.70 | 1.20 | 1.12–1.28 |
| **By number of affected first-degree relatives** |  |  |  |  |
| 0 | 2544 | 2857.48 | 0.89 | 0.85–0.92 |
| 1 | 412 | 371.93 | 1.10 | 1.00–1.22 |
| 2 | 35 | 23.02 | 1.52 | 1.09–2.11 |
| **Combined model - 10-year risk: (No first-degree family history)** |  |  |  |  |
| Quintile 1 (lowest) | 124 | 171.16 | 0.72 | 0.60–0.86 |
| Quintile 2 | 329 | 383.60 | 0.85 | 0.77–0.95 |
| Quintile 3 | 465 | 592.75 | 0.78 | 0.71–0.86 |
| Quintile 4 | 664 | 767.75 | 0.86 | 0.80–0.93 |
| Quintile 5 (highest) | 962 | 942.22 | 1.02 | 0.95–1.08 |
| Top 10% | 546 | 499.22 | 1.09 | 1.00–1.19 |
| Top 5% | 305 | 259.97 | 1.17 | 1.04–1.31 |
| Bottom 10% | 36 | 63.30 | 0.56 | 0.41–0.78 |
| Bottom 5% | 14 | 26.90 | 0.52 | 0.30–0.87 |
| **Combined model – Full lifetime risk: (no first-degree family history)** |  |  |  |  |
| Quintile 1 (lowest) | 333 | 479.16 | 0.69 | 0.62–0.77 |
| Quintile 2 | 438 | 541.21 | 0.81 | 0.73–0.88 |
| Quintile 3 | 494 | 579.08 | 0.85 | 0.78–0.93 |
| Quintile 4 | 547 | 609.60 | 0.89 | 0.82–0.97 |
| Quintile 5 (highest) | 732 | 648.44 | 1.12 | 1.05–1.21 |
| Top 10% | 408 | 330.10 | 1.23 | 1.12–1.36 |
| Top 5% | 214 | 167.53 | 1.27 | 1.11–1.46 |
| Bottom 10% | 151 | 229.02 | 0.66 | 0.56–0.77 |
| Bottom 5% | 79 | 111.42 | 0.71 | 0.56–0.88 |

SIR was calculated based on number of cases observed and expected using sex-specific UK population rates of colorectal cancer incidence rates, calculated for the entire eligible UK Biobank cohort or by family history status and stratified by full lifetime and 10-year risk
categories for the combined model (except for first-degree relative, where categories of number of first-degree relative were used). Abbreviations: O = observed, E = expected.

### Model performance

For full lifetime risk, the AUC for the combined model was 0.673 (95% CI 0.664–0.682 and the AUC for the family history model was 0.666 (95% CI 0.657–0.675). For 10-year risk, the AUC of the combined model was 0.674 (95% CI 0.665–0.683) and the AUC of the family history model was 0.668 (95% CI 0.659–0.677). The difference between the model fits was significant (10-year risk: χ^2^=7.16, df=1, p=0.0075; full lifetime risk: χ^2^=7.42, df=1, p=0.0065). The 10-year risk combined model was slightly under-dispersed (dispersion coefficient 1.08, 95% CI 1.07–1.09), while the full life risk combined model was considerably under-dispersed (dispersion coefficient 1.84, 95% CI 1.83–1.86). Further supporting this data, when we analysed the observed and expected ratio (SIR) using 10-year follow-up data, we noticed an overestimation of risk for both the family history model (SIR=0.94, 95% CI 0.91–0.98) and the combined model (SIR=0.95, 95% CI 0.91–0.98), compared with the population incidence data.

### Risk stratification

We investigated the risk distributions of the family history model (Fig 1A, C) and the combined model (Fig 1B, D) using the entire eligible UK Biobank cohort. Fig 1A shows the full lifetime risk distribution for the family history model, where there are six possible categories (0, 1 and 2+ affected first-degree relatives by gender) (median=0.073, inter-quartile range=0.027, min=0.053, max=0.188). Fig 1B shows the full lifetime risk distribution for the combined model (median=0.057, inter-quartile range=0.042, min=0.011, max=0.688). Fig 1C and Fig 1D show the 10-year risk distribution for the family history model (median=0.010, inter-quartile range=0.009, min=0.001, max=0.055) and the combined model (median=0.008, inter-quartile range=0.009, min=0.0004, max=0.251) respectively.

**Fig 1:**
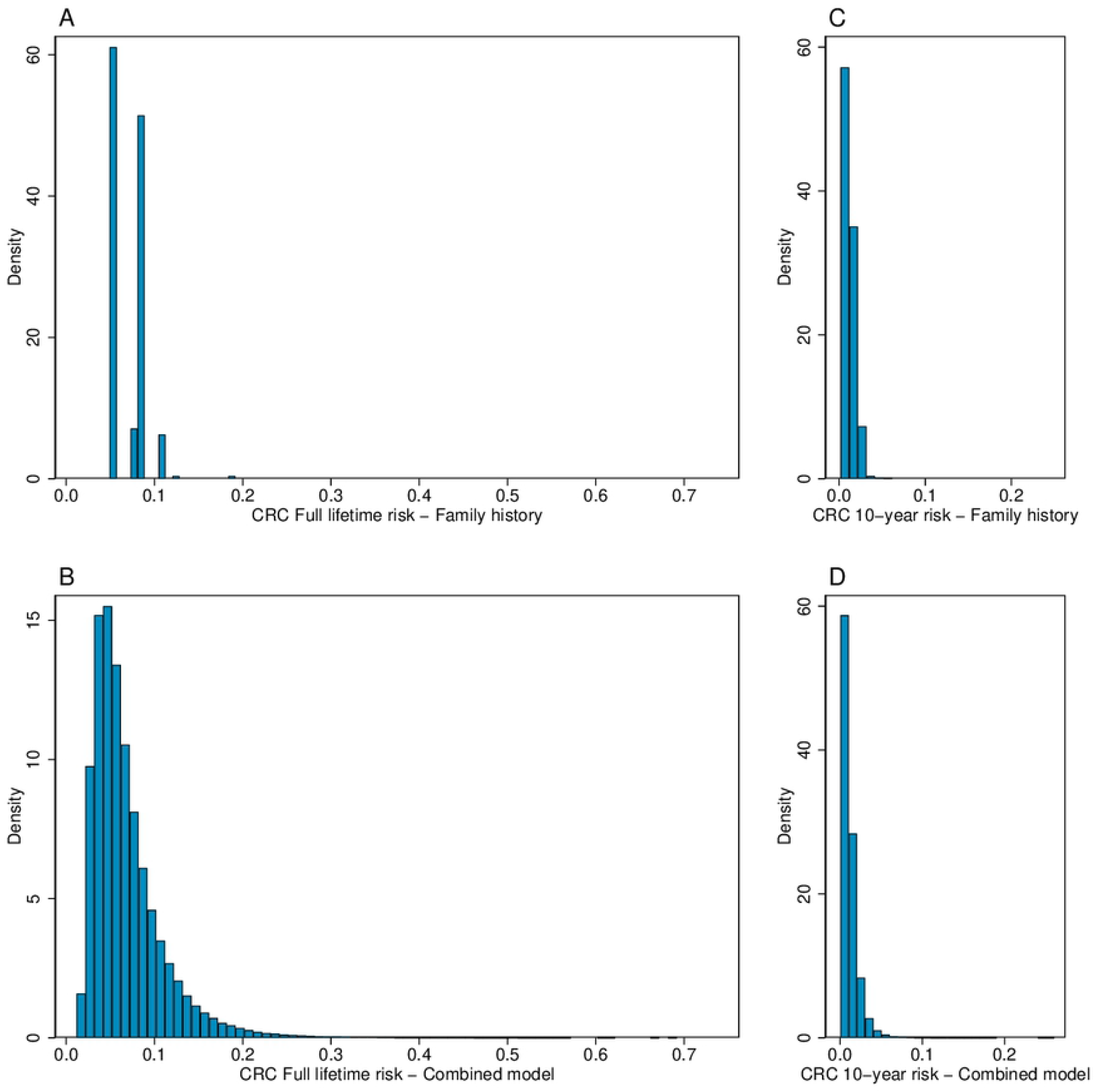
Risk distribution plots for the eligible UK Biobank participants. Plots show the Full lifetime risk distribution for a model with family history only (A) and the combined model (B), and 10-year risk for the family history model (C) and the combined model (D).

The SIRs by quintiles of full lifetime risk and 10-year risk for the combined model are shown in Table 3 and Fig 2. We observed an increase in risk gradient between full lifetime risk categories; persons in the top quintile of risk have ~35% higher colorectal cancer incidence than those in the middle quintile and ~53% higher colorectal cancer incidence than those in the bottom quintile. The 10-year risk quintile gradient was less than the full lifetime risk gradient, but showed the same trend. To compare risk stratification of persons with a family history to those without, we also broke down the SIR analysis by number of affected first-degree relatives and for people without any family history. We observed that the top quintile, decile and 95^th^ percentile (for participants without family history) have similar risk values, compared to someone with 1 affected first-degree relative. Also, the risk for people with 2 affected first-degree relatives overlaps with the top risk categories (due to the large confidence interval) (Fig 2). Although the range of SIR is diminished in the 10-year risk graph, the trend is still visible (Fig 2D).

**Fig 2:**
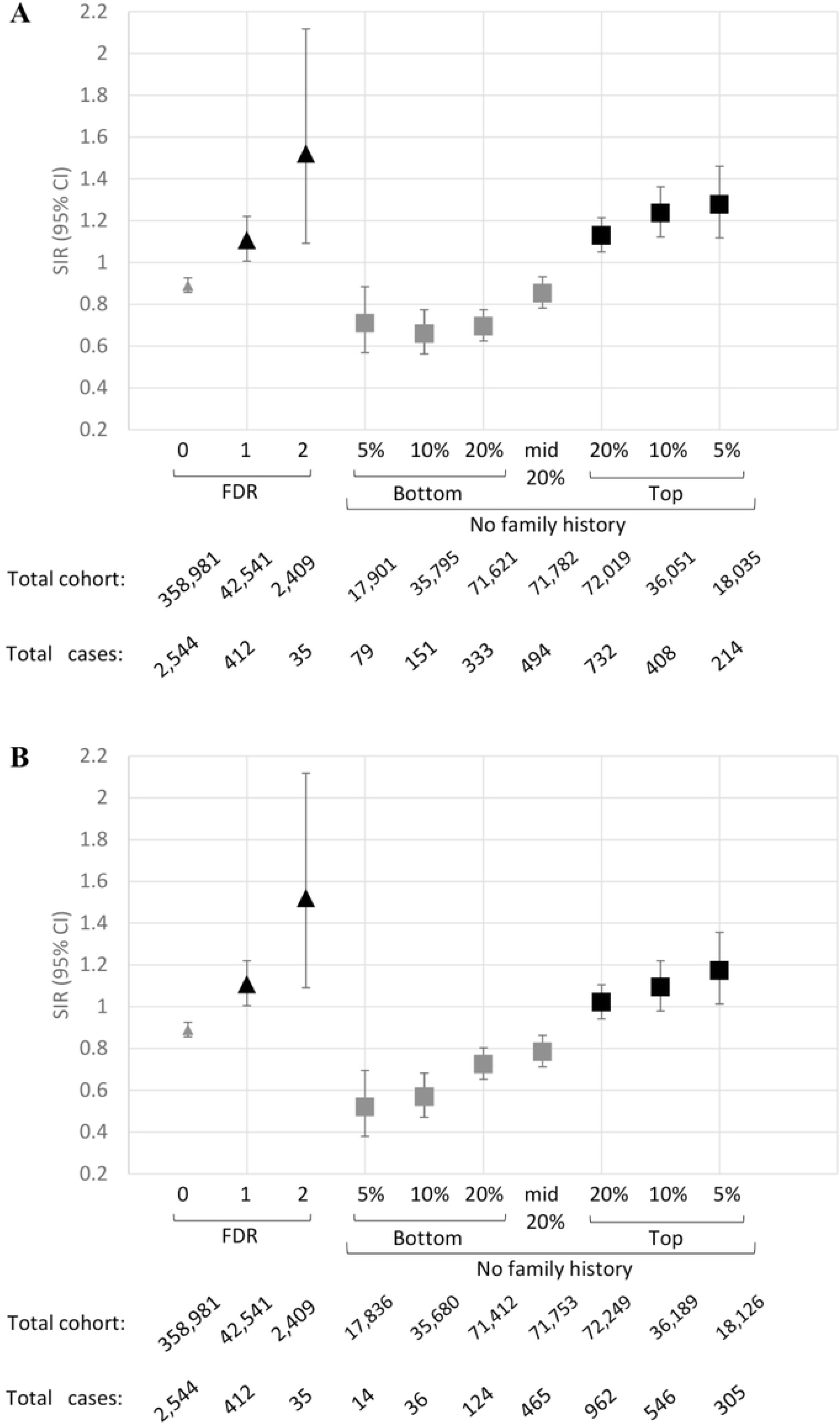
Comparison of the standardised incidence ratios (SIR) for different subgroups. SIR values were generated based on number of cases observed and expected using sex-specific UK population incidences for the number of affected first-degree relatives (FDR) vs the combined model for people without a family history. SIR values were plotted against number of affected first-degree relatives in comparison with full lifetime (A) and 10-year (B) risk categories for participants without family history.

## Discussion

Colorectal cancer is a major public health issue worldwide, with high incidence in many westernised countries (34), in addition to increasing incidence for young adults (35, 36). Several modalities for early detection exist including colonoscopy and faecal occult blood testing. Evident in long-term trends available from US Surveillance, Epidemiology, and End Results data, prevention of advanced colorectal cancer is feasible with screening programs based on colonoscopy as opposed to faecal occult blood testing. In comparison with UK and Australian data, US colorectal incidences are below their Western counterparts and this could be due to differences in screening programs. But every program comes at a cost (colonoscopy versus faecal occult blood testing, for example). The US approach (by the American Cancer Society) (37) to rising incidence in young adults is to lower colonoscopy screening age to 45 - an approach that will surely be more effective at detecting early onset disease, but at a far greater cost and an increased risk of over screening thousands of adults. Furthermore, Ladabaum et al. (38) provided evidence that screening compliance would be a more efficacious approach to reduce colorectal incidence and death. The National Colorectal Roundtable announced in 2018 the goal of achieving 80% colorectal screening participation in every community in the US. Building on that compliance goal, and keeping early onset disease in mind, novel risk stratification approaches can only improve screening outcomes by enabling a focus on at-risk persons.

The majority of colorectal cancer cases do not have monogenic (Lynch syndrome and familial adenomatous polyposis) causes, but have (39) multifactorial causes due to genetic, environmental and lifestyle factors (1). Risk stratification of the general population will assist in identifying those at higher risk and enable the implementation of targeted screening and risk reduction for this group. Currently, screening decisions in the general population are based on age and family history (in UK, USA, and Australia), and recommendations for early screening are based on the number (and age at diagnosis) of affected first-degree and second-degree relative(s) (as determined by each country’s medical bodies) (40–42). Basing screening decisions on family history alone has its caveats, including incorrect reporting of cases in relatives due to lack of knowledge of the cancer diagnosis, or of the site of the cancer (43, 44). However, as the vast majority of colorectal cancer cases do not have a first-degree relative (3, 4, 45), current screening guidelines do not accurately identify persons above population average risk thresholds. There are many other factors involved in the risk colorectal cancer, including genetic, environmental and lifestyle factors that, if measured and taken into account, can more accurately identify where people are with respect to risk-based screening thresholds. Given that there is the potential for more than 40% of colorectal cancer cases to be prevented by behavioural modification, risk-stratification based on non-modifiable risk factors (like family history and polygenic risk) could allow for pre-emptive screening and, importantly, cost-effective risk-reduction options. Notably, the potential benefits of a so-called “healthy lifestyle” on colorectal cancer incidence appears to be evident across all polygenic risk categories (46, 47).

In this study, we evaluated how much a PRS based on 45 SNPs (19) improves colorectal cancer risk prediction when added to a simple family history model. By confirming the performance of a PRS originally constructed using a cohort enriched for family history (19), we have therefore demonstrated the clinical validity of this risk measure for the general population. We have shown that adding a PRS to a model that includes only family history results in modestly improved discriminatory performance. We have also shown that the variance of the risk distribution of the combined model is much greater than that of the family history alone model, Fig 1). As a consequence, for the UK Biobank participants, using the combined model there were more than 29,000 (~17% of) males with no affected first-degree relative but a full lifetime risk scores greater or equal to ~11% (the family history model risk score of a male with one affected first-degree relative in the UK Biobank). There were more than 34,000 (~17% of) females with no affected first-degree relative with full lifetime risk scores (combined model) greater or equal to ~7.3% (the family history model risk score of a female with one affected first-degree relative in the UK Biobank). In agreement with previous data (48–50), using a PRS we are able to identify 72,019 participants with an increased risk equivalent to having an affected first-degree relative. Importantly, the combined model captures the crucial components of non-modifiable colorectal cancer risk.

In summary, we have found that stratifying colorectal cancer risk by including a PRS with first-degree family history results in an improved risk prediction compared with using family history alone for a sample that mirrors the general population, for which <15% had a family history. This is in agreement with previous studies that have also shown that a PRS adds substantial value in colorectal cancer risk stratification and explains a sizeable excess risk of colorectal cancer, independent of family history (17, 48). Our new data strengthens the argument for clinical application of polygenic risk assessment in the general population, and especially for those without a family history, and supports the expansion of current recommendations that focus only on family history and age as the main criteria for screening.

Better colorectal cancer risk stratification in the general population will improve identification of at-risk individuals. A significant finding of our work is that 20% of participants based on PRS have a similar full lifetime risk of colorectal cancer as the ~11% identified solely by a first-degree family history, and therefore should thus be assessed with the same importance. Reinforcing the importance of the polygenic risk score for assessing risk is the recent finding that four of the SNPs included in our PRS (rs12241008, rs2423279, rs3184504, and rs961253) have been shown to be associated with increasing adenoma count at colonoscopy (51). Adenoma count is not only an indicator of risk itself but is a measure of colorectal cancer development. Identification of at-risk individuals based on a PRS-integrated model will allow for the improved screening and thus removal of such lesions prior to malignant transformation. Furthermore, there is evidence that the PRS association is stronger for proximal compared with distal disease (52) suggesting that risk assessment could help inform endoscopists’ colonoscopy procedural plan, such as a slightly slower withdrawal time (53).

From a health economic perspective, the model used in the present study, which incorporates only non-modifiable risk factors, exceeds the benchmark discrimination threshold (AUC ≥ 0.67) at which risk stratified colorectal cancer screening is thought to become cost effective (54). Future iterations of the combined model to include additional risk factors should only improve the calibration and discrimination and consequently improve the clinical utility of such a tool for colorectal cancer screening uptake, compliance and screening cessation, and post-polypectomy follow-up.

### Conclusion

The practical clinical benefit of a risk assessment model that combines PRS and family history is to identify adults who are at an increased risk of colorectal cancer, sufficient to qualify for supplemental screening recommendations who would not otherwise be identified because they do not have a family history, or do not have a strong enough family history to meet screening thresholds.

### Study limitations

Our study has several limitations. First, our model is under-dispersed, both for the 10-year risk and full lifetime risk models, although this might be corrected once we update the model with additional risk factors. Furthermore, the model only takes into account first-degree relatives, and we do not break down the risk of first-degree relative by consultand’s age or age at diagnosis of the first-degree relative. Given that familial risk, and the PRS associations, depend on these ages (55), there will be some underestimation of risks for young adults and some overestimation of risk for the majority of adults with mild family history—such as a first-degree relative diagnosed at 70 or older. In two recent meta-analysis, the overall colorectal cancer risk associated with family history was found to be lower than previously reported, suggesting we are likely over-estimating familial risk in older adults (27, 56). Because of the substantial environmental contribution to colorectal cancer, there remains unaccounted, modifiable risk not captured by this combined model. Calibration of the model could be improved by increasing the number of susceptibility SNPs and adding further clinical risk factors in the models including smoking history, alcohol and processed meat consumption and BMI. Secondly, the UK Biobank recruited only between the ages of 40 69 years. There were few incident cases in participants in their 40s. This affected our ability to confirm published evidence (17, 48, 57), suggesting superior clinical utility in PRS to help detect early-onset colorectal cancer before age 50 years; we observed a not significant trend in the expected direction for the few young age at diagnosis cases (S2 Table). Furthermore, a recently published study (58) has identified, using exome data, 76 participants in the UK biobank who are potential Lynch syndrome carriers, 17 of whom are cases. Although these are small proportions of the cohort, they could still bias our results, causing an underestimation in some of our standard incidence ratio estimates. To investigate this, we excluded participants from the analysis based on the published pathogenic variants initially identified (58) and compared the SIR results to the original dataset. We found no difference in comparison with the original analyses, as the majority of these potential Lynch syndrome participants didn’t pass our eligibility criteria for the analysis (Table 1), resulting in only two Lynch syndrome cases in the final dataset. Furthermore, as shown in Table 3, SIR estimates from the entire cohort were lower than expected (by ~8%), suggesting the UK biobank cohort is “healthier” with respect to colorectal cancer risk than the general population. “Healthy lifestyle” is associated with colorectal cancer incidence regardless of PRS, and because we do not yet incorporate modifiable risk factors, our model is not accounting for those who are at high risk based on the PRS but are at low risk based on modifiable risk factors, and vice versa. This could also affect the performance of our model in stratifying colorectal cancer risk categories. Additionally, we do not account for risk differences for those participants who underwent bowel screening. Therefore, we could be overestimating short-term risk for those who have had bowel screening. Ten-year risk scores are meant to assess short-term risk of being diagnosed with colorectal cancer and would be more efficacious for the general population if modifiable risk factors were incorporated. This current model incorporates non-modifiable risk factors and is best suited for determining baseline colorectal cancer risk without the consideration of highly modifiable risk factors attributed to colorectal cancer (59). Finally, we are aware of the population-specific limitations of this study which was restricted to white, Northern-European population. While there is evidence that many susceptibility SNPs are consistent in the strengths and direction of their associations across ethnicities (60–62), there are ethnic specific-loci and variants that have yet to be incorporated into this model.

### Future directions

To improve the model calibration, we plan to perform future analysis using additional colorectal cancer susceptibility risk SNPs and create an expanded combined model with additional risk predictors to produce a more comprehensive colorectal cancer risk assessment tool, applicable across multiple ethnicities. Improvement and validation of the predictive ability of such a colorectal cancer risk assessment tool will facilitate implementation and ultimately hopefully adoption into routine clinical care.

## Acknowledgments

We wish to thank Mr Lawrence Whiting for his invaluable expertise in the management of large data files from the UK Biobank.

